# A Retrospective Analysis of Career Outcomes in Neuroscience

**DOI:** 10.1101/2024.02.01.578220

**Authors:** Lauren E. Ullrich, John R. Ogawa, Michelle D. Jones-London

**Affiliations:** NINDS

## Abstract

What factors are associated with different career outcomes among biomedical PhDs? Much of the research to-date has focused on drivers of interest in (and intention to pursue) various careers, especially during graduate school, but fewer studies have investigated the ultimate career outcomes of participants. Even less is known about what factors matter most for groups historically underrepresented in the US STEM workforce, such as for women, some racial and ethnic groups, and persons with disabilities (National Center for Science and Engineering Statistics (NCSES), 2021a). This study reports a new analysis of data from 781 PhD neuroscientists that were obtained from a retrospective survey (reported in Ullrich et al. (2021)) to investigate the factors that influence the career sector in which neuroscience PhDs are employed, and whether there were group differences according to social identity. We find evidence of academia as a “default path” for incoming PhD students, but interest in different careers changes gradually over time. Those who remained in academia had greater acceptance of the structural aspects of academic careers, such as the promotion and tenure process, and greater faculty support during their postdoctoral training. Conversely, prioritizing monetary compensation and wanting varied work were associated with not being in academia, while a strong interest in research was positively associated with being in non-academic research. Somewhat surprisingly, there were few interactions with gender, and no interactions with underrepresentation status. Our findings also underscore the role of advisors, networking, and personal relationships in securing employment in STEM.

## INTRODUCTION

What factors drive career outcomes among biomedical PhDs? Much of the research to-date has focused on drivers of interest in (and intention to pursue) various careers, especially during graduate school. Academic faculty careers are often framed as the “default career” during biomedical PhD training (Sauermann & Roach, 2012), a framing evident in qualitative analysis of free-text responses to this survey (Ebrahimi et al., 2022), high levels of interest at start of PhD (Gibbs et al., 2014; Ullrich et al., 2021), and “excessively optimistic” beliefs about the chance of obtaining an academic career (Ganguli et al., 2022; Puljak & Sharif, 2009). However, interest in research-focused faculty careers decreases during PhD training, a trend amplified in women and members of US-based historically underrepresented (UR) racial and ethnic groups (C N Fuhrmann et al., 2011; Gibbs et al., 2014; Golde & Dore, 2004; Roach & Sauermann, 2017; Sauermann & Roach, 2012; but see Wood et al., 2020).

Research on the development of career interest in STEM PhD students has shown that it is a multifactorial process that evolves over time. It is influenced by factors such as confidence in one’s ability as a researcher (self-efficacy), social and intellectual feelings of belonging, interactions with one’s advisor, perceptions of conflicting time demands of work and family, and institutional climate or culture (Curry & DeBoer, 2020; Estrada et al., 2011, 2019; Gazley et al., 2014; Gibbs & Griffin, 2013; Hayter & Parker, 2018; Kahn & Ginther, 2017).

Fewer research studies have investigated factors related to the ultimate choice of career (as opposed to interest in or intention to pursue a career), but since career interest likely plays a role in that choice, similar factors to those mentioned above are likely to play roles. Additional pressures such as changes in family obligations, low wages, availability of jobs, and even external biases during the job search may also influence the ultimate sector in which someone is employed (Quadlin 2018; Way et al. 2016; Rivera 2017; Bloch et al. 2015).

In addition, it is possible that whatever factors affect ultimate career choice might affect members of different demographic groups (e.g., women and/or UR minority groups) in different ways, resulting in the unequal representation of certain groups in STEM careers reported by the National Center for Science and Engineering Statistics NCSES) (2021a). This unequal representation holds true in academic research specifically—and equal representation decreases at every progressive step along the academic track. Although underrepresented racial/ethnic groups comprise 38% of the US population, only 16% of the PhD recipient pool (National Center for Science and Engineering Statistics (NCSES), 2021b), 19% of medical school graduates (Association of American Medical Colleges, 2023a), 10% of current assistant professors, and 7% of tenured faculty at U.S. medical schools according to publicly available data (Association of American Medical Colleges, 2023b). Demographic differences are also apparent in career sectors across STEM, such as government and industry (Kahn & Ginther, 2017; Mathur et al., 2018). Data from the National Science Foundation show several differences in employment outcomes by demographic variables. In academia, women biology PhD holders are more likely than men to be employed at educational institutions other than 4-year institutions (such as community colleges, two-year colleges, and technical and vocational schools) and private non-profits (National Center for Science and Engineering Statistics (NCSES), 2021a). Black and Hispanic biology PhD holders are more likely than non-Hispanic white and Asian biology PhD holders to be employed in government and educational institutions other than 4-year institutions (National Center for Science and Engineering Statistics (NCSES), 2021).

We previously reported (Ullrich et al., 2021) the results of a survey of recent neuroscience PhD graduates in which we found repeated evidence that individual preferences about careers in general, and academic careers specifically, predict current career interest. Recapitulating trends found in the broader STEM workforce (Gibbs et al. 2015; Gibbs et al. 2014; Sinche 2016; Layton et al. 2016; Lambert et al. 2020), these findings were moderated by social identity (a combination of gender and membership in an UR minority group) and experiences in graduate school and postdoctoral training. A subset of that sample had completed their training and entered the workforce. In this study we investigated which factors, including career interest, were associated with participants’ actual career choices, namely the job sector of their current position. We consider social identity, career interest, experiences in graduate school and postdoctoral training, personal characteristics, and measures of research experience and productivity and examine their relation to career choice. It is important to note that the relations that we report are not causal, but associative, since this is an exploratory retrospective study and all variables were measured at the same time.

## MATERIALS AND METHODS

### Sample

The study population was composed of: 1) current graduate students or recent doctoral recipients (calendar year 2008 or later), 2) who were US citizens or permanent residents, and 3) had applied for individual NINDS funding or have been appointed to institutional NINDS training (T32) or research education grants (R25). Participants were identified within the National Institutes of Health (NIH) database and invited by email to complete the survey during summer 2017. The survey received 2,675 responses, representing approximately 36% of identified individuals and 45% of opened emails. After data cleaning there were 2,242 complete, eligible, and unique response. Additional information on recruitment and data collection is described in (Ullrich et al., 2021). As determined by the NIH Office of Human Subjects Research, federal regulations for the protection of human subjects did not apply to this activity.

### Definitions and Sample Refinement

Several criteria were applied to the 2,242 responses to refine the sample for analysis. Since social identity, namely gender and race/ethnicity, were of primary interest for this article, responses that did not include that information were excluded, leaving 2,065 responses. Second, only participants who answered that their current position was “Professional in the field” were included, leaving 866 participants. This restriction meant that participants had completed their training and had chosen at least an initial career (i.e., they were not current graduate students or postdoctoral fellows). Finally, only those in science or science-related careers were retained (i.e., not unemployed or working outside of science), which led to a final total of 781 participants.

Participants included all who answered either “male” or “female” and may include transgender respondents who identify as either a man or woman. Only two participants indicated “other” and wrote in a response for gender; they were not included in the analysis due to small numbers. Respondents from white and/or Asian backgrounds are referred to as well-represented (WR), while respondents from American Indian/Alaska Native, Black/African American, Hispanic/Latino, and/or Native Hawaiian/Pacific Islander backgrounds are referred to as underrepresented (UR), according to the NSF definition (National Science Foundation, 2015). Moving forward, this is referred to as representation status. Disability status was collected, but persons with a disability made up less than 3% of the final sample, so were not included as a separate analysis group because of the small sample size.

### Survey

The survey was a 57-question instrument administered at a single point in time (Ullrich et al., 2021; Appendix 1). The questions were iteratively developed by synthesizing from several sources, conducting cognitive testing interviews, and refining language where necessary (Ullrich et al., 2021). The survey asked about respondents’ current position; demographics; career interest; experiences in graduate school and postdoctoral training; personal characteristics; and objective measures of research experience and productivity.

Respondents were asked to categorize their current position into one of the following career groups: Academic position, research focus (includes physician-scientist); academic position, teaching focus; non-academic research (e.g., research in industry, biotech, or government settings); science-related non-research (e.g., science outreach, communication, policy, advocacy, or administration); and other, non-science-related careers (Gibbs et al., 2014; National Institutes of Health, 2012). Non-science-related careers had fewer than 60 cases and were not included in the analyses for this paper since they did not represent a specific career sector. Respondents were also asked about their social identity, specifically gender and race/ethnicity, and other demographic information.

Respondents were asked to rate their interest in pursuing each of the above career pathways at three time points: the start of their PhD program, the end of their PhD program, and currently. Interest was measured on a 4-point Likert-type scale where 1 = no interest, 2 = low interest, 3 = moderate interest, and 4 = strong interest.

Experiences in training included: various aspects of their relationship with their primary training advisor during graduate and postdoctoral training (5-point scale from “very negative” to “very positive”); sources and helpfulness of support and career advice during the graduate and postdoctoral training (4 point scale from “no guidance provided” to “very helpful”); and feelings of social and intellectual belonging to lab/research group and department/program during graduate and postdoctoral training (5-point scale from “strongly disagree” to “strongly agree”).

Personal characteristics included: confidence in one’s potential as an independent researcher (measured on a 5-point agreement scale where 1 was “strongly disagree” and 5 was “strongly agree”); aspects of the career or work environment most important to the respondent (choose up to top 5); and features of academia that increase or decrease desire to become a faculty member (5-point scale from “greatly decrease” to “greatly increase”).

Objective measures of research experience and productivity included: years of research prior to PhD program, total years of research, years to complete PhD, total time in postdoctoral training, years since PhD completion, support by NIH prior to the PhD program, first-author publication rate (first-authored publications/total years performing research), time to PhD completion, and undergraduate or doctoral degree from a top 50 research university (as measured by research and development expenditures, National Science Board, 2016).

### Analysis

This work was designed to investigate whether individual characteristics, experiences during training, and attitudes and preferences around careers are associated with the type of current position participants held. The outcome variable was nominal (type of current position) which had the following categories: research-focused academic, teaching-focused academic, non-academic research, and science-related non-research.

#### General Notes

All data analyses were conducted using version 4.0.5 of the R program (R Core Team, 2018). Individual packages are cited in text when referenced. Prior to any analyses, all continuous variables were visually checked for outliers by plotting and comparing to similar curves, and any outliers were recoded to the largest/smallest value that fit the visual curve (cap method). Using this method, four observations were capped. All interactions were evaluated in the context of component main effects and all lower-level interactions and continuous variables were centered.

Definitions for small, medium and large effect sizes for this article were generally taken from Cohen (1988): mean differences used *d* (small (s) ≥ 0.2, medium (m) ≥ 0.5, large (l) ≥ 0.8), and correlation coefficients and individual regression coefficients used *r* (s ≥ 0.1, m ≥ 0.3, l ≥ 0.6). Finally, for odds ratios we used (rounded) Cohen’s cutoffs (s ≥ 1.5, m ≥ 2.5, l ≥ 4.5). Significance was indicated by adjusted p values < .05.

#### Data Reduction

Data reduction was performed for several constructs to reduce multicollinearity, multiple comparisons, and Type I error. We used factor analysis to reduce these constructs (e.g., relationship with advisor) into latent factors.

For each analysis the number of factors to extract was ascertained using BIC scores computed through the VSS function from the psych package in R (Revelle, 2019). Then, the fa function (also from the psych package) was used to compute maximum-likelihood solutions, with oblique rotation performed using the “promax” option. Twenty-six questions were reduced to 9 factor variables.

#### Changes in Career Interest Ratings Over Time: Repeated Measures MANOVAs

Changes over time in participants’ reported interest in the four career types that might have differed by their current position were investigated through repeated measures MANOVAs. Both time (“Time:” T1, T2, T3) and interest in the different types of careers (“Type:” research-focused academia, teaching-focused academia, non-academic research, and scientific non-research) were within-subjects dependent variables. Between-subjects independent variables were current position, gender, and UR status. First an omnibus repeated measures (RM) MANOVA containing all dependent and independent variables was computed using RM from the MANOVA.RM package (Friedrich, Konietschke & Pauly, 2021). P values were computed using 1,000 iterations of RM’s Wild Bootstrap option for ATS (ANOVA-Type Statistic), and then adjusted using Benjamini and Hochberg’s (1995) procedure (see below). Since the Time by Type by Current Position interaction was significant, along with several lower-order interactions, four follow-up Time by Type repeated measures MANOVAs were conducted within each level of Current Position, again using RM. False Discovery Rate (FDR) was controlled using Benjamini and Hochberg’s (1995) procedure, which was applied to each analysis using the “BH” option on the mt.rawp2adjp function of R’s Multtest package (Gentleman et al., 2005).

#### Gender and Representation Status Differences in Current Position and Explanatory Variables: Chi-squared Test, Logistic Regression, Multinomial Logistic Regression, and ANOVA

Gender and representation status differences in current position and the set of 56 explanatory variables were investigated using analysis techniques that matched the nature of the dependent variables (here current position and the explanatory variables): a contingency table and chi-squared for current position, logistic regression for the dichotomous explanatory variables, multinomial logistic regression for the multinomial explanatory variables, and ANOVA for continuous explanatory variables.

Gender and representation status differences on current position, a multinomial categorial variable, were investigated by computing Pearson’s chi-squared test of goodness-of-fit on the contingency table formed by crossing social identity (gender and representation status) with current position (Table 2). Follow-up chi-squared statistics were computed for each sub-table (within social identity type). For sub-tables with significant differences, standardized cell residuals (Z-scores) were computed for each cell.

Gender and representation status differences on explanatory variables were investigated through three different procedures, depending on the nature of the explanatory variable. For all three sets of analyses, each analysis used the explanatory variables as dependent variables, and Gender, Representation Status, and their interaction as the independent variables. False Discovery Rate (FDR) was controlled using the procedure from Benjamini & Hochberg (1995), which was applied to each analysis using the “BH” option on the mt.rawp2adjp function of R’s Multtest package (Pollard et al., 2005)

Gender and representation status differences on the 18 dichotomous explanatory variables were investigated through logistic regressions computed using glm from the base package in R with family = “binomial.” Gender and Representation Status differences on the three multinomial explanatory variables were investigated through multinomial logistic regressions computed using multinom from the nnet package in R (Venables & Ripley, 2002). Statistics for individual terms were computed by successively contrasting statistics from the full model to statistics from three sub-models that each had a different term removed using the ANOVA test for model comparison, so FDR adjustment was not possible. Finally, gender and representation status differences on the 35 continuous explanatory variables were investigated through ANOVAs computed using aov from the base package in R.

#### Two-Step Logistic Regressions Investigating Factors Associated with Current Position

Three two-step analyses were performed, one for each of the decisions presented in the results section: 1) academia vs. non-academia, 2) research-focused academia vs. teaching-focused academia, and 3) non-academic research vs. science-related non-research. These branching dichotomies were chosen based on the qualitative analysis of free text responses, which indicated that careers in academia were the “default path” for the participants (Ebrahimi et al., 2022). Each of the three analyses followed the same procedures (outlined below)—only the dependent variables changed between analyses.

Since we had 56 possible independent (explanatory) variables and 2 possible moderators (gender and representation status) which might be associated with each of the three dependent variables (decisions 1 through 3), it was imperative to reduce the set of independent variables and moderation terms to only those which might be related to each of the dependent variables in a multivariate context to control both Type I error and multicollinearity. We chose to utilize a strategy first introduced by Efron et al. (2004) as the LARS-OLS Hybrid and proved mathematically by Belloni & Chernozhukov (2013), in which a large set of possible independent variables is reduced using penalized regression (lasso in our case) techniques, and the resulting set of independent variables are analyzed with ordinary least squares (OLS) methods.

As an initial step in these analyses, we imputed values for any missing variables using the mice package in R (van Buuren & Groothuis-Oudshoorn, 2011)

The first step, lasso logistic regression, was conducted using the glinternet package in R (2021; Michael Lim & Hastie, 2015). The glinternet.cv procedure allows the computation of group-lasso, in which groups of variables can be linked together such that their coefficients are always estimated together, an important requirement for testing interactions. It can require strong hierarchy, such that interactions are always estimated within the context of their main effects. Each of the 56 explanatory variables was crossed with gender, representation status, and the gender by representation status interaction and considered as a main effect without interaction such that the final group of variables entered into the glinternet.cv equation had 227 terms.

The gender by representation status interaction was computed separately and entered as if it was a single variable instead of an interaction term since no group-lasso procedure had the capability to handle 3-way interactions without including all possible interactions among independent variables, a predictor pool that would have been too large for this study. This may have led to some two-way interactions or Gender and Representation Status main effects not being part of the 3-way interaction groups, producing inflated coefficients for some of the 3-way interactions. However, the only major consequence of our workaround would be that these 3-way interactions were non-significant in the second step OLS logistic regressions and would have effectively reduced degrees of freedom, thus making the OLS logistic regressions more conservative.

The glinternet.cv package also performs cross-validation, in which successive random subsets of the sample are analyzed leaving a specified portion of the sample (1/5^th^ in our case) left out for “out of box” testing of each model (for an overview, see James et al., 2013)

Given our interest in inference, we used a method for choosing the penalty term, lambda, that maximized coefficient size, within the generally accepted bounds of lambda.min and lambda.1se, by computing the mean coefficient size for the equation generated by each lambda between and including lambda.min and lambda.1se. We then chose the lambda that maximized coefficient size as our penalty term (Hastie et al., 2009).

Once we had chosen lambdas for each of the three lasso logistic regressions, we computed the final equations using those lambda values. These equations provided the final sets of variables that would be used in the second-step OLS logistic regressions.

The three second-step OLS logistic regressions took all variables and interactions that had non-zero coefficients in their respective first step lasso logistic regression as their independent variables and the variable representing the decision of interest for that analysis as the dependent variable. OLS logistic regressions were performed using the glm procedure with family = “binomial” from base R. FDR was controlled in these analyses using the procedure from Benjamini & Hochberg (1995) as outlined above.

#### Contingency Table Analyses Investigating Path to Current Position

The contingency table partitioned the sample into a matrix that had answers to how the participants found their current positions in rows and current position type in columns. An omnibus chi-squared statistic was computed for the table to assess any association between the two variables using chisq.test in base R. We used the “simulate.p.value” option in chisq.test to require 2,000 Monte Carlo draws to arrive at a robust p value because some cells may have had low counts.

Because the omnibus chi-squared test was significant, we continued with follow-up analyses to determine which rows had significantly different distributions from the overall distribution using chisq.test to compare each row to the overall row sums. Again, the “simulate.p.value” option was used to offset any bias from low cell counts. The p values for the 6 follow-up tests were adjusted for FDR using the procedure from Benjamini & Hochberg (1995), which was applied using the “BH” option in the adjust.p procedure in base R.

Finally, in rows that were found to be significantly different from the overall row distribution for the table, individual cells were compared to all other cells from the same row to identify groups of cells that were similar. We wanted to examine each method of finding a position (in rows) to find out which of the current position types (in columns) used it most often and which used it least often. Cell counts or row percentages would have been biased for this purpose by differences in the frequencies of current position types. Therefore, we compared the column percentage that was associated with each cell to the column percentages of the other cells in that row using chi-squared tests of independent proportions (prop.test in base R). FDR was controlled for these follow-up cell comparisons by adjusting the p values for the 6 follow-up tests for each row using the procedure from Benjamini & Hochberg (1995), again applied using the “BH” option in the adjust.p procedure in base R.

## RESULTS

### Sample Demographics

Table 1 presents basic demographic information about the 781 PhD neuroscientists in the sample. All reported that they were “professionals in the workplace,” indicating that their training was complete. Participant information includes gender (54% women, n=422) and representation (WR/UR) status (16% UR, n=123). Current Position, the main outcome for the study, is also reported in Table 1. Slightly more than half, 51%, reported that they were in research-focused academic positions; 13% in teaching-focused academic positions; 17% in non-academic research positions; and 19% in science-related non-research positions. In addition, 45% held a PhD in neuroscience, the rest were in biological or health-related fields. The median years since PhD for the sample was six, and 74% of respondents had been a postdoctoral fellow. Respondents from 193 PhD institutions in 46 states, the District of Columbia, and Puerto Rico were represented.

**Table 1.**
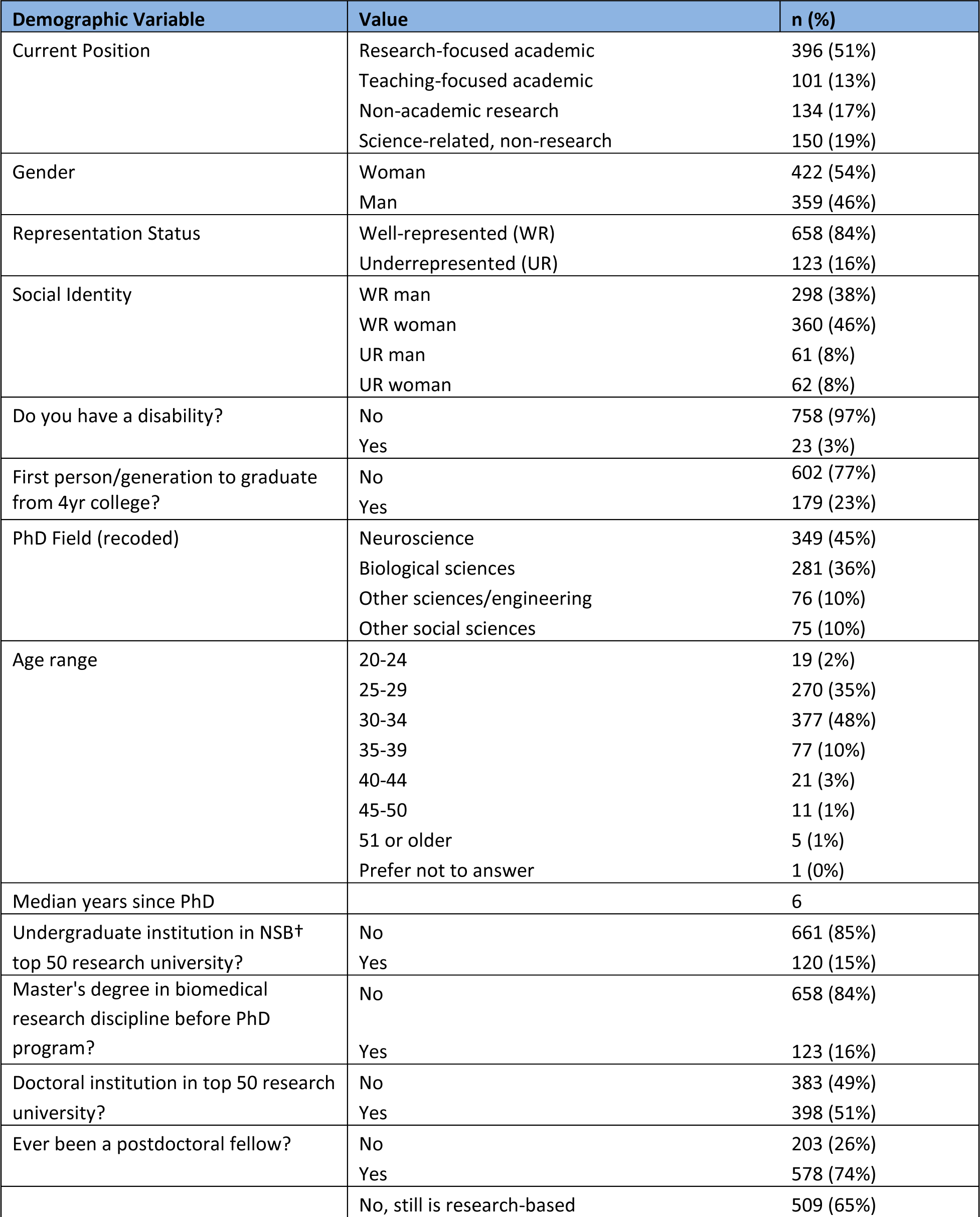
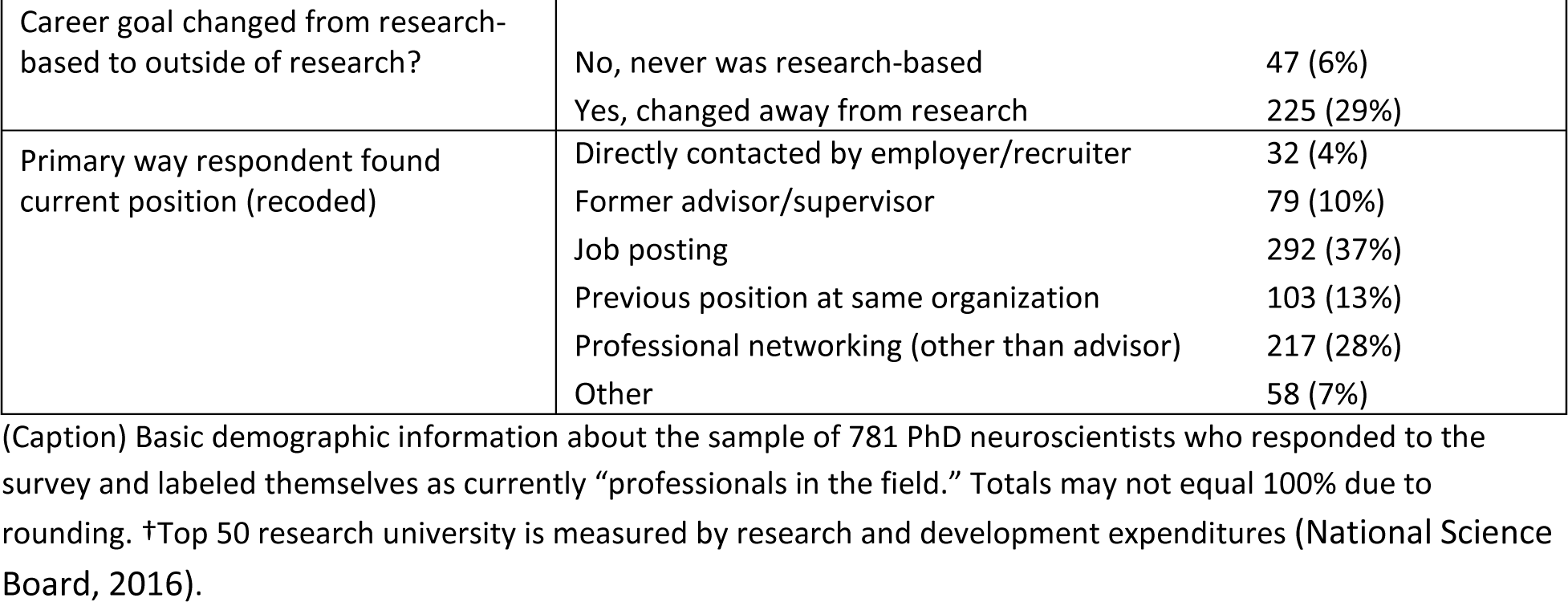
Study sample characteristics.

The sample is not a random sample but does have similar sample demographics as other studies (Gibbs et al., 2014). Due to the source for contact information (NIH’s database and internet searches) and the fact that respondents had to have either applied for NIH funding or been supported by NIH funding, there is likely an overrepresentation of people in academic positions.

### Approach

After characterizing demographic information, we first investigated the relationship between career interest and the ultimate choice of career. We looked for changes in how respondents who ended up in each of the career sectors reported their career interest over time. We then investigated whether women and UR respondents reported different experiences in graduate school and postdoctoral training, personal characteristics, and objective measures of research experience and productivity when compared to men and WR respondents, respectively. Next, we investigated which of these factors were associated with participants’ current positions. Finally, we investigated whether participants’ reports of how they found their current positions differed by the type of position.

### Changes in Career Interest Over Time Related to Current Position

First, we investigated whether participants’ reported interest in the four career types at three time points (start of PhD, end of PhD, and current) were related to their current positions. For example, did participants who are now in research-focused academic positions have different patterns of career interest over time than participants who are now in non-academic research positions? To answer this question, we split participants by their current position and analyzed the change over time in their report of interest levels in various career types (see Figure 1 and Extended Data Figure 1-1 to 1-4).

**Figure 1.**
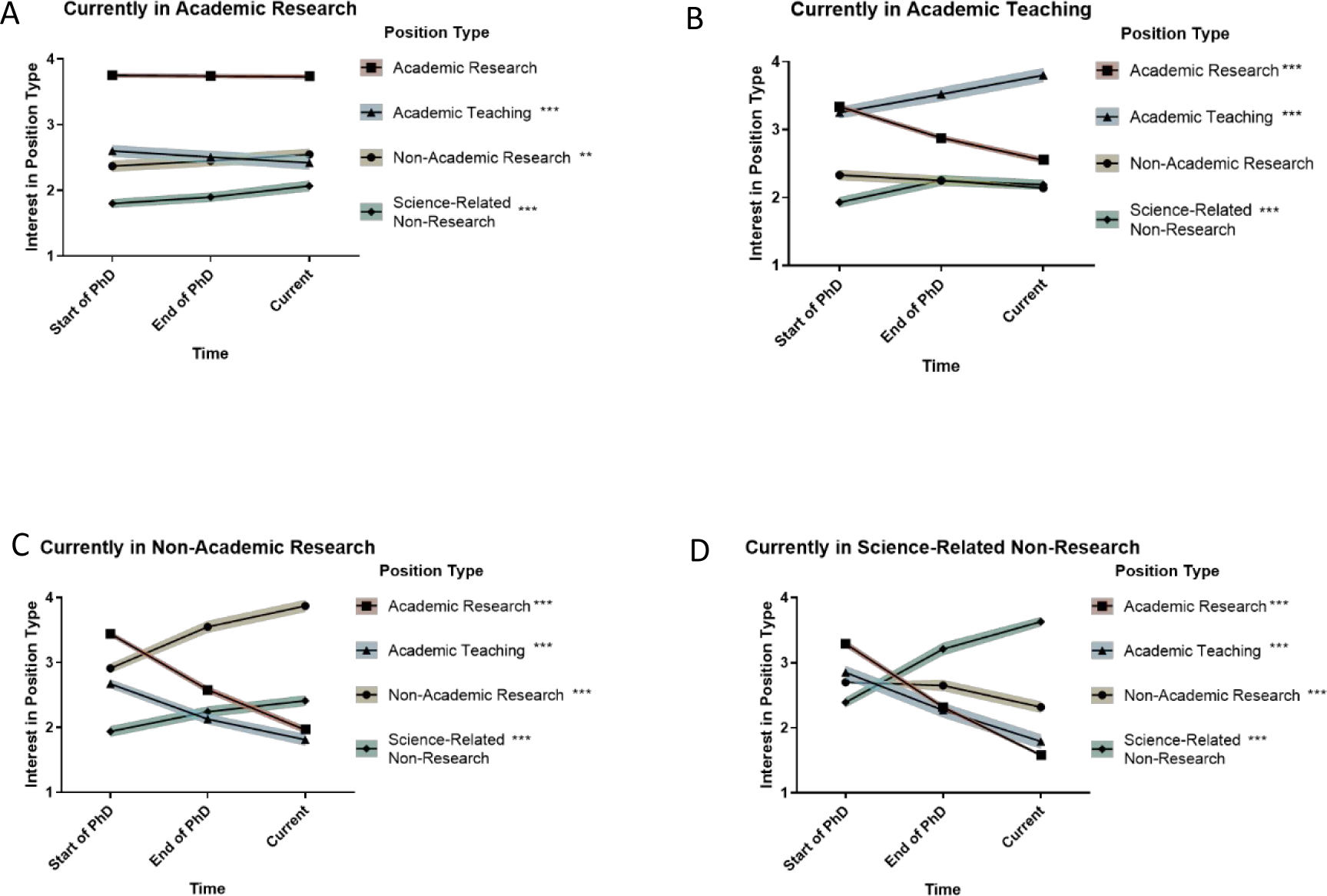
Career interest has solidified by end of PhD. Graphs show changes in career interests over time, split by current position. Mean responses of PhD neuroscientists in four different career paths who were asked to rate their level of interest in four different career paths at three time points: start of PhD, end of PhD, and current, on a 4-point scale (where 1 represents “no interest” and 4 represents “strong interest”). Repeated measures MANOVA showed that mean retrospectively reported interest in all four career types changed over time for participants in all career types except academic research-focused respondents’ interest in academic research (A) and academic teaching-focused respondents’ interest in non-academic research (B) (Extended Data Figure 1-1 to Figure 1-4; standard error around the mean is indicated by colored shading; **p < 0.01, ***p < 0.001; for M, SD, and effect size detail see Extended Data Figure 1-4).

The preliminary omnibus repeated measures MANOVA investigating within-subjects dependent variables time and career interest rating by between-subjects independent variables current position, gender, and representation status (see Extended Data Figure 1-1) found one significant 3-way interaction: current position by time by career interest rating. Follow-up time by career interest rating repeated measures MANOVAs, split by type of current position, found significant time by career interest rating interactions for each type of current position (see Extended Data Figure 1-2). Findings were followed-up by further repeated measures (time) ANOVAs split by current position and career interest ratings (see Extended Data Figure 1-3). These findings, along with means, are presented in Figure 1 and in Extended Data Figure 1-4.

For each of the four current positions, the mean retrospectively reported interest rating changed over time for almost all career types, as demonstrated by the interest ratings of almost all types of careers, split by participants’ current positions, showing a significant within-subjects time effect (see Figure 1 and Extended Data Figure 1-4).

The first finding of note is that for those in research-focused academic positions (Figure 1A), the career type with the highest mean interest rating did not change over time—it was always research-focused academic positions. In addition, while their interest in teaching-focused academic positions decreased over time, their interest in non-academic research and science-related non-research increased slightly (but significantly) over time.

For participants in the three other types of positions (teaching-focused academic, non-academic research, and scientific non-research) the patterns of interest were different from those of research-focused academics (Figure 1B-D). For all three of the groups, the career type with the highest level of interest at the start of PhD was research-focused academia. At the end of PhD, participants report that their highest mean level of interest matched their current positions (e.g., participants who were currently non-academic researchers report the highest mean interest in non-academic research at the end of their PhD), often at a higher mean level than their initial interest in research-focused academia at the start of PhD. At the time of the survey (“current interest”), the career type with the highest mean level of interest remained the same as the participants’ current positions, but with an additional increase over their ratings at the end of PhD. In addition, interest in teaching-focused academic positions decreased over time for everyone except those currently in teaching-focused academic positions. In contrast, interest in science-related non-research positions increased for those in all four types of positions.

These patterns demonstrate that, in general, although most participants reported that they were interested in research-focused academia at the start of their PhD training (T1), by the end of their PhD training (T2) those who would end up in different career types reported that their interests had shifted to their current career type. In addition, their current interest (T3) only reinforces this shift by demonstrating an increased level of interest in their current career type.

Science-related non-researchers showed the greatest changes in interest over time, with the greatest decrease in interest in research-focused academic positions and the greatest increase in interest in their current career type. Non-academic researchers showed the second greatest changes in interest over time.

### Differences in Current Position by Gender and Representation Status

As a prelude to analyses examining social identity and other factors that related to current position, we investigated whether there were any differences in participants’ current positions related to their social identity. Table 2 presents the results of a contingency table analysis crossing current position by social identity. The omnibus chi-squared test of independence was significant (Χ^2^(9)=70.1, p<0.001), so follow-up goodness-of-fit tests (testing against the overall distribution of current positions) were performed for each social identity’s distribution of current positions. Three of the four follow-ups were significant, indicating that the current position distribution for those social identities were different from the overall distribution of current positions. Finally, adjusted standardized residuals were computed for each of the three significant follow-up distributions, allowing us to determine which cells were contributing the most to the significant chi-squared statistics. These residuals, and equivalent p-values based on Z-score cutoffs, are displayed in the cells of Table 2.

**Table 2.** Current position by gender and representation status.

| Current Position | Social Identity |  |  |  | TOTAL |
| --- | --- | --- | --- | --- | --- |
|  | UR Women*** | WR Women** | UR Men | WR Men*** |  |
| Research-focused academic | 22 (35%)* | 157 (44%)* | 30 (49%) | 187 (63%***) | 396 (51%) |
| Teaching-focused academic | 9 (14%)* | 56 (16%) | 8 (13%) | 28 (9%) | 101 (13%) |
| Non-academic research | 6 (10%) | 54 (15%) | 17 (28%) | 57 (19%) | 134 (17%) |
| Science-related, non-research | 25 (40%***) | 93 (26%**) | 6 (10%) | 26 (9%***) | 150 (19%) |
| **TOTAL** | 62 (100%) | 360 (100%) | 61 (100%) | 298 (100%) | 781 (100%) |

(Caption) Proportion of “professionals in the field” respondents in each of the four “current positions” by gender and representation status. UR = underrepresented, WR = well represented. Significance level indicators (\* $p < 0.05$ , \*\* $p < 0.01$ , \*\*\* $p < 0.001$ ) in column headings show subset chi-squared goodness-of-fit tests comparing the distribution of current positions for that social identity and the overall distribution of current positions. Significance level indicators in individual cells show Z-score tests for adjusted standardized residuals computed for that social identity group alone.

UR women were much more likely to be in science-related non-research positions, and less likely to be in research-focused academic positions than the group as a whole. WR women were more likely to be in science-related non-research positions and less likely to be in research-focused academic positions than the group as a whole. Conversely, WR men were much more likely to be in research-focused academic positions, and less likely to be in science-related non-research positions than the group as a whole.

### Differences in Explanatory Variables by Gender and Representation Status

Next, we asked whether graduate school and postdoctoral training experiences, personal characteristics, and objective measures differed by social identity in our sample. Significant findings were followed-up by examining differences either in means or slopes for subsamples defined by whichever was significant of gender, representation status, or their interaction (Figure 2 and Extended Data Figure 2-1 to Figure 2-3).

**Figure 2.**
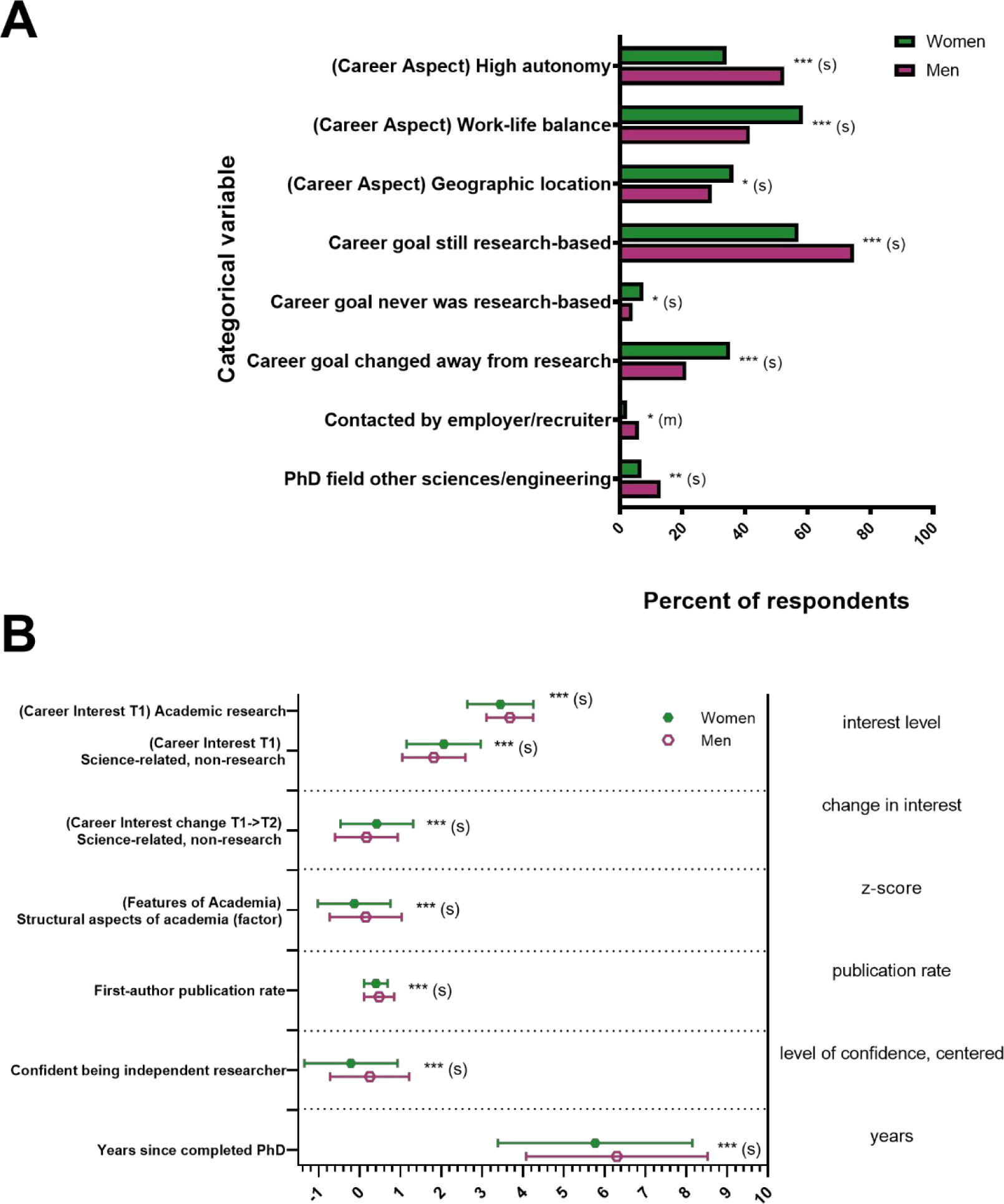
Gender differences among responses to variables capturing experiences, personal characteristics, and objective measures. Significance levels from F statistics (ANOVA, reported in Extended Data Figure 2-1 to Figure 2-3) comparing the means for women and men for each variable were all significant at p < 0.05, at least. A) Categorical variables. Responses on the X-axis were the percent of respondents in each group who indicated either that the response was important to them or agreed with the response. B) Interval variables. Responses on the X-axis are indicated to the right of the graph: interest levels in a given career (1 to 4, 4 being very interested); T2 interest minus T1 interest for change in interest; z-score for “structural aspects of academia” factor scores; total publications/years of research for publication rate; level of confidence (centered −3.05 to 0.95, 0.95 being most confident) for confidence in being an independent researcher; and years since completed PhD. Effect sizes are labeled when they reach at least “small” size. (s) = small effect size, (m) = medium effect size, *p < 0.05, **p < 0.01, ***p < 0.001. Error bars indicate standard deviation.

#### Gender

We found significant differences in training experiences, personal characteristics, and objective metrics between men and women (significant results in Figure 2, all results in Extended Data Figure 2-1 to Figure 2-3; all results in Figure 2 were significantly different and had at least a small effect size).

Women were less likely to have received their PhDs in the “other sciences/engineering” category of fields than men were. Women in our sample also report they completed their PhD more recently than men, and have lower publication rates than men. In addition, women’s current ratings of their confidence in their potential to be independent researchers were lower than men.

Women also had different ratings than men of the importance of various aspects of careers – women rated “autonomy” less important, and “work/life balance” and “geographic location” more important than men. For the factors assessing whether different features of academia increased or decreased their interest in academia, women reported that the “structural aspects of academia” factor decreased their interest in academia more than it decreased men’s interest. Women were also approached directly for their current positions by employers or recruiters less than men; this was the only result that had a medium effect size.

Women’s reports of their interest in different career types at the start of their graduate training also differed from men’s recall of their interests. At T1, women were less interested in academic positions with a research focus and more interested in science-related non-research positions than men. In addition, women’s interest in science-related non-research positions increased more over the course of their graduate training than men’s interest did. Finally, women were more likely to respond that their career goals had changed away from research positions or had never been research positions compared to men.

#### Representation Status

WR and UR respondents differed significantly on objective measures (significant results in Figure 3, all results in Extended Data Figure 2-1 to 2-3; all results in Figure 3 were significantly different and had at least a small effect size). We found that UR respondents were more likely to have been supported by NIH NRSA awards fewer times before their current position than WR respondents; UR respondents took longer to obtain their PhDs; and UR respondents reported lower publication rates than WR respondents. Additionally, UR respondents were twice as likely as WR respondents to have been the first person or in the first generation of their family to graduate from a 4-year college or university and were less likely to have received their PhDs in the “Other sciences/engineering” category of fields than WR respondents; both had a medium effect size.

**Figure 3.**
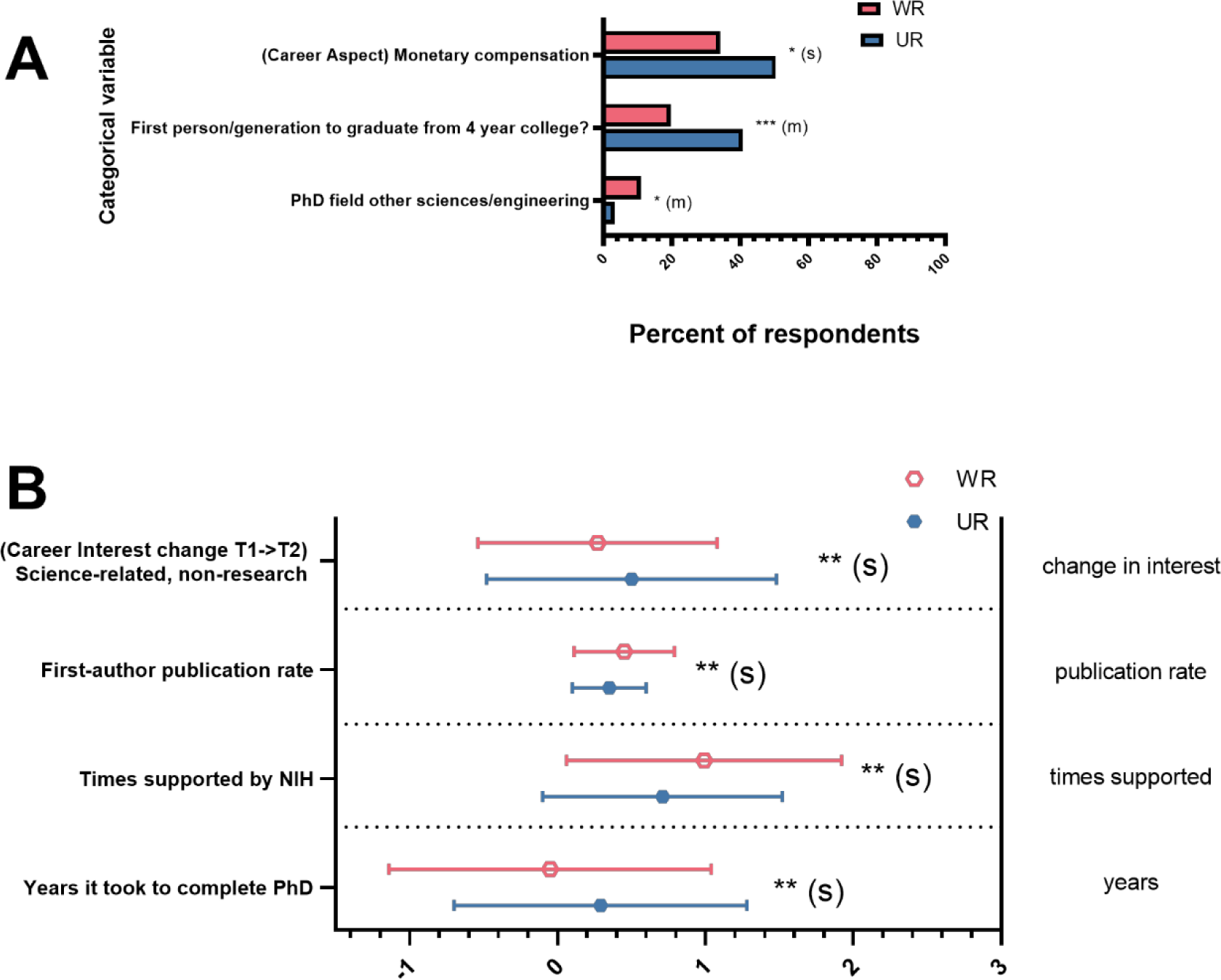
Differences between WR and UR responses to variables capturing experiences, personal characteristics, and objective measures. Significance levels from F statistics (ANOVA, reported in Extended Data Figure 2-1 to 2-3) comparing the means for WR and UR responses for each variable were all significant at p < 0.05, at least. A) Categorical variables. Responses on the X-axis were the percent of respondents in each group who indicated either that the response was important to them or agreed with the response. B) Interval variables. Responses on the X-axis were T2 interest minus T1 interest for change in interest; total publications/years of research for publication rate; times supported by NIH; and years (centered) for years it took to complete PhD. Effect sizes are labeled when they reach at least “small” size. (s) = small effect size, *p < 0.05, **p < 0.01, ***p < 0.001. Error bars indicate standard deviation.

We also found differences between UR and WR respondents in personal characteristics. UR respondents selected “monetary compensation” as an important aspect of career choice more often than WR respondents. Finally, UR respondent interest in science-related non-research positions increased more over the course of their graduate training than did WR respondent interest, even though the level of initial interest in these positions did not differ between UR and WR respondents (t(161)=-1.28, p=n.s.).

There were no significant interactions between gender and representation status for any of the explanatory variables (Extended Data Figure 2-1 to Figure 2-3).

### Associations with Current Position

Our main question was: which factors were associated with the type of position that respondents were in currently? The analyses that addressed this question used the same broad array of explanatory variables as examined above: experiences in graduate and postdoctoral training, personal characteristics, and objective measures. In these analyses, however, they were used as independent variables along with gender and UR status, and all of the interactions amongst them.

We analyzed our data according to the two levels of choices that mirror the decision-making process that recent PhDs and postdocs use to decide what type of jobs to pursue (this strategy of analyzing hierarchies of discrete choices has long been employed in economics through decision tree analysis, e.g., see Magee (1964)). The first choice is whether to stay in academia (Ebrahimi et al., 2022; Gaughan & Robin, 2004; Helbing et al., 1998; Puljak & Sharif, 2009). Then, for academics, the decision is between faculty positions that are research-focused and those that are teaching-focused. For non-academics, the decision is between research positions (i.e., research in industry or government labs) and those that are science-based, but not research positions (i.e., in policy, non-profits or government).

We chose this 2-level approach because we posit that predicting all four careers in one analysis with the entire sample might wash out similar but different associations between the outcome and the predictor variables that would be clearer when examined only for the sub-groups for whom the decision mattered. For example, in the whole sample an interest in research might predict membership in both the non-academic research group and the research-focused academic group, which would cause any variance associated with group differences to drop out of the analysis. By separating the sample into two analyses on two different sub-samples, namely

1. Determining characteristics that differentiated research vs. teaching positions in only those respondents who were in academic positions, and,
2. Determining characteristics that differentiated research vs. non-research positions in only those respondents who were in non-academic positions,

we can capture the utility of an interest in research in differentiating the four types of positions.

#### Factors Associated with Being in Academic vs. Non-Academic Positions

First, we examined the associations between explanatory variables and whether respondents’ current positions were in academia or not. The details of the 2-step process we used in arriving at a final logistic regression predicting academic vs. non-academic current positions are described in the Analyses section. The first step, lasso logistic regression investigating factors associated with being in academic vs. non-academic positions (Extended Data Figure 4-1), provided the list of 28 explanatory/independent variables that were entered into the second step OLS logistic regression. The second step logistic regression equation (Figure 4, Table 3) was performed on data from all 781 participants. The equation had a significant log-likelihood test, correctly predicting whether 89% of participants were in academia or were not in academia. Thirteen predictors were significant and had sufficient effect size to report.

**Figure 4.**
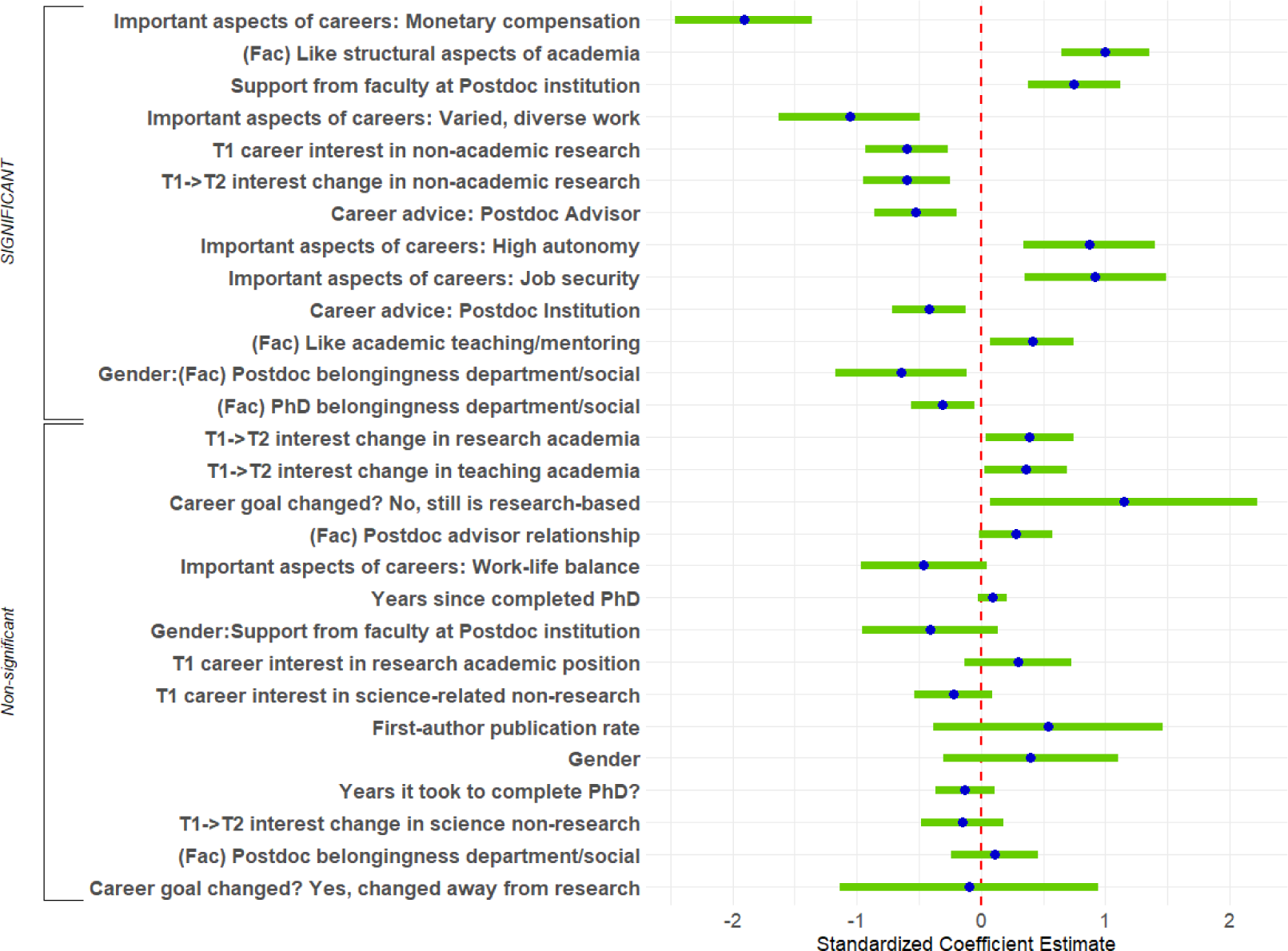
Logistic Regression Predicting Academic vs. Not Academic Positions. Standardized regression coefficients and error bars for logistic regression predicting whether respondents were in academic or non-academic positions. The dependent variable was the binary indicator of whether respondents’ positions were academic. Independent variables included career interest and change in career interest, experiences during PhD training and postdoctoral training, personal characteristics, objective measures, and interactions with gender and representation status. Factors are indicated by (f) and coded (not raw) values are indicated by (c). The entire equation was significant at p<0.001 and accurately predicted 89% of respondents in the analysis.

**Table 3.**
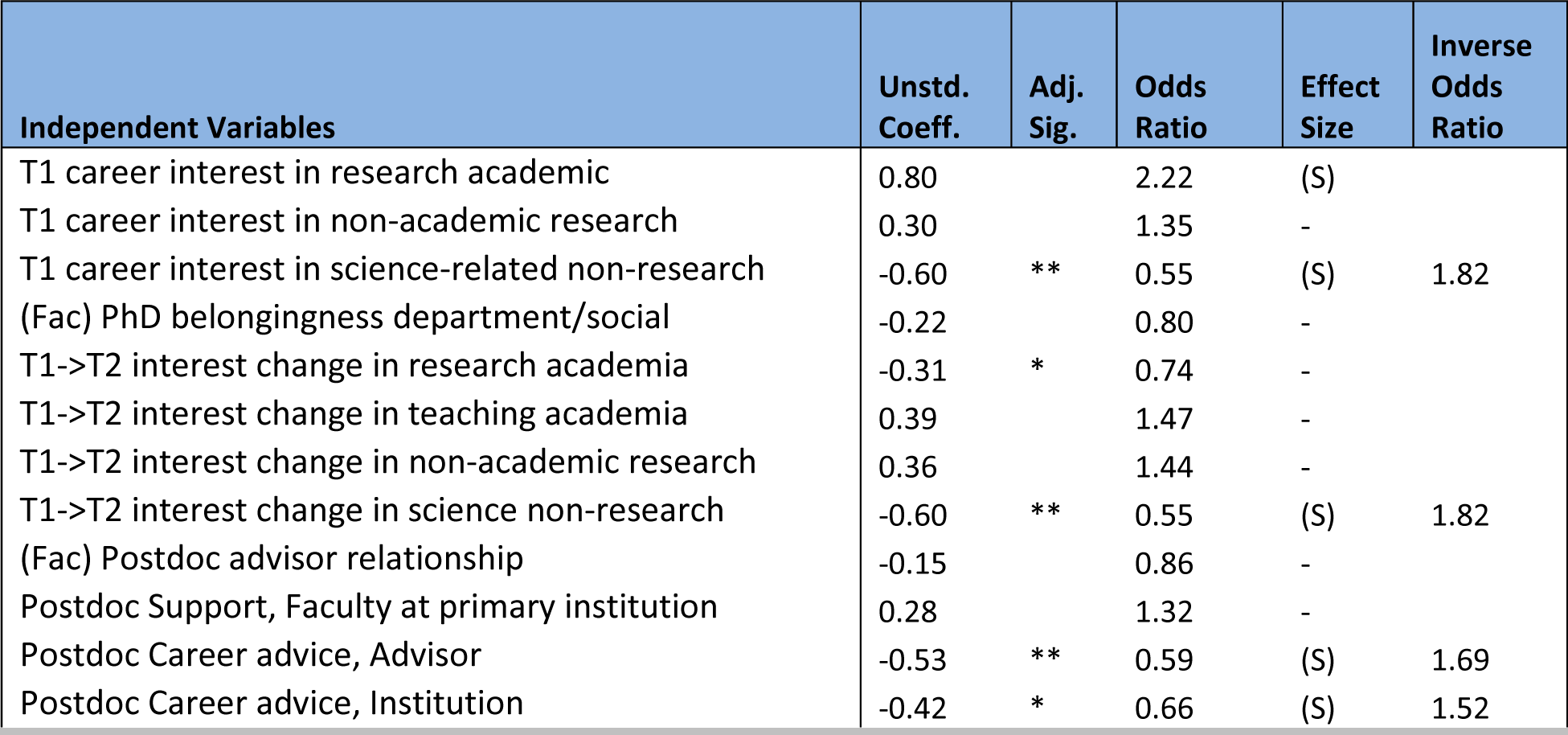
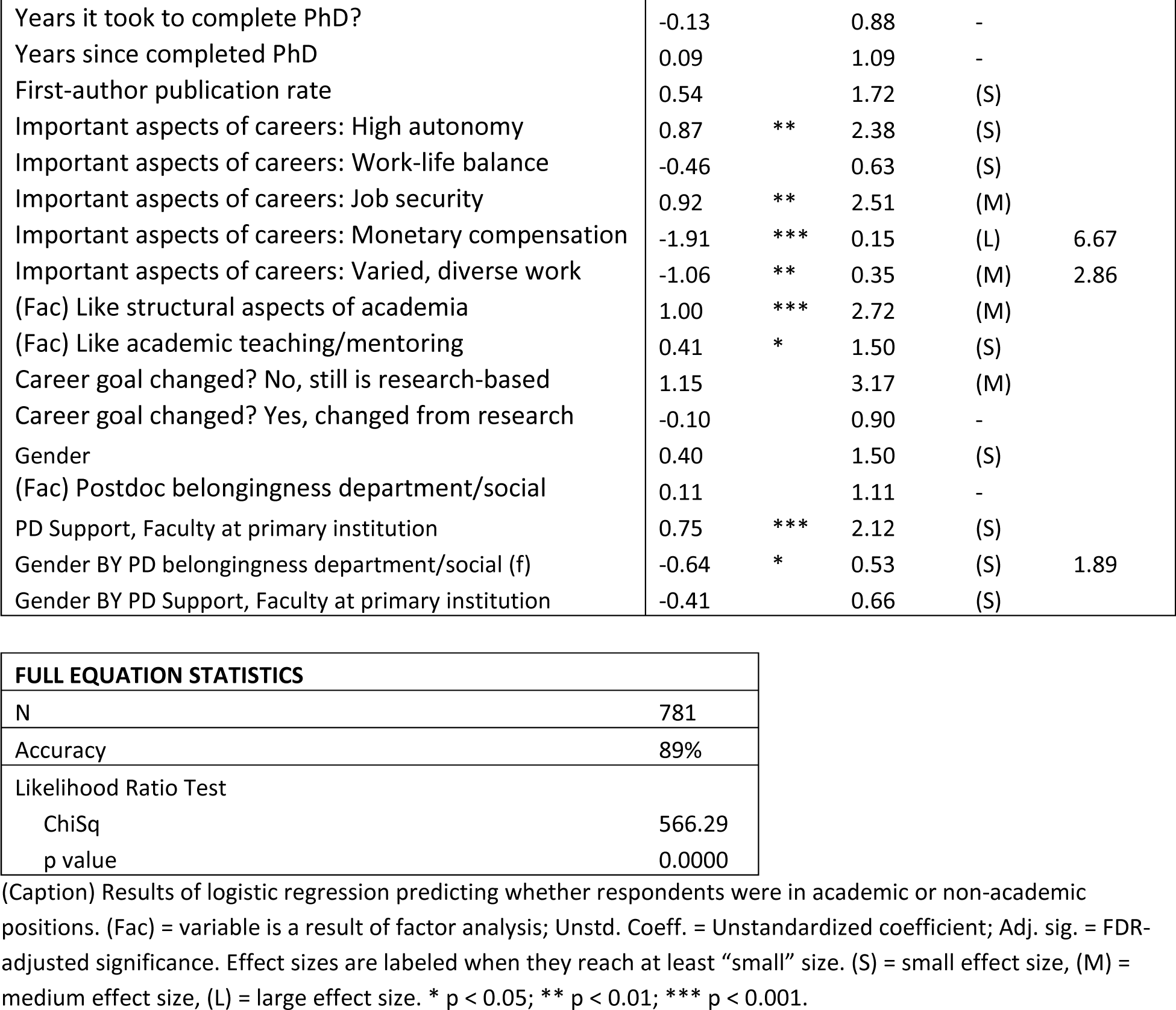
Logistic regression predicting career in academia vs. not academia.

Several personal preferences were associated with whether participants were in academia. On the one hand, wanting job security and high autonomy were positively associated with being in academia. On the other hand, feeling that monetary compensation was an important aspect of one’s career was strongly positively associated with not being in academia (large effect size), as was wanting varied/diverse work (medium effect size). Participants’ feelings about particular features of academia were also associated with being in academia, such as feeling positive about the structural aspects of academia (job market, promotion and tenure) and feeling positive about teaching/mentoring.

Participants’ experiences as postdocs were also associated with being in academia, albeit in sometimes unintuitive ways. First, feeling that faculty support at their primary institutions was helpful was positively associated with being in academia. Rating career advice from advisors or institutions as helpful, however, was positively associated with not being in academia. Finally, feeling like one belonged to the intellectual/social community of their postdoc departments affected women and men differently (an interaction). For women, feelings of departmental belongingness were significantly positively associated with being in academia (Intellectual: χ^2^(4)=23.87, p<0.0001 and Social: χ^2^(4)=13.12, p<0.0107). For men, feelings of belongingness were not related to being in academia.

Perhaps unsurprisingly, participants’ interest in non-academic research positions was associated with being in non-academic positions. High interest in non-academic research at the start of graduate training and increases in interest in non-academic research during graduate training were also both associated with being in non-academic positions.

Other single variables were not reported because they were either significant but their effect sizes did not meet the threshold for reporting or had reportable effect sizes but were non-significant. No interactions with representation status remained significant in this analysis.

#### Factors Associated with Being in Research-focused vs. Teaching-focused Academic Positions

Next, we examined the associations between explanatory variables and respondents’ type of academic position: research- or teaching-focused. The first step lasso logistic regression investigated factors associated with being in a research-focused academic position vs. a teaching-focused academic positions. Extended Data Figure 5-1 provided the list of explanatory/independent variables that were entered into the second step OLS logistic regression. The second step logistic regression equation (Figure 5, Table 4) was performed on data from the 497 participants who were in academia. The equation had a significant log-likelihood test, predicting whether 90% of participants were in research-focused academic positions or teaching-focused academic positions. Seven predictors were significant and had sufficient effect size to report.

**Figure 5.**
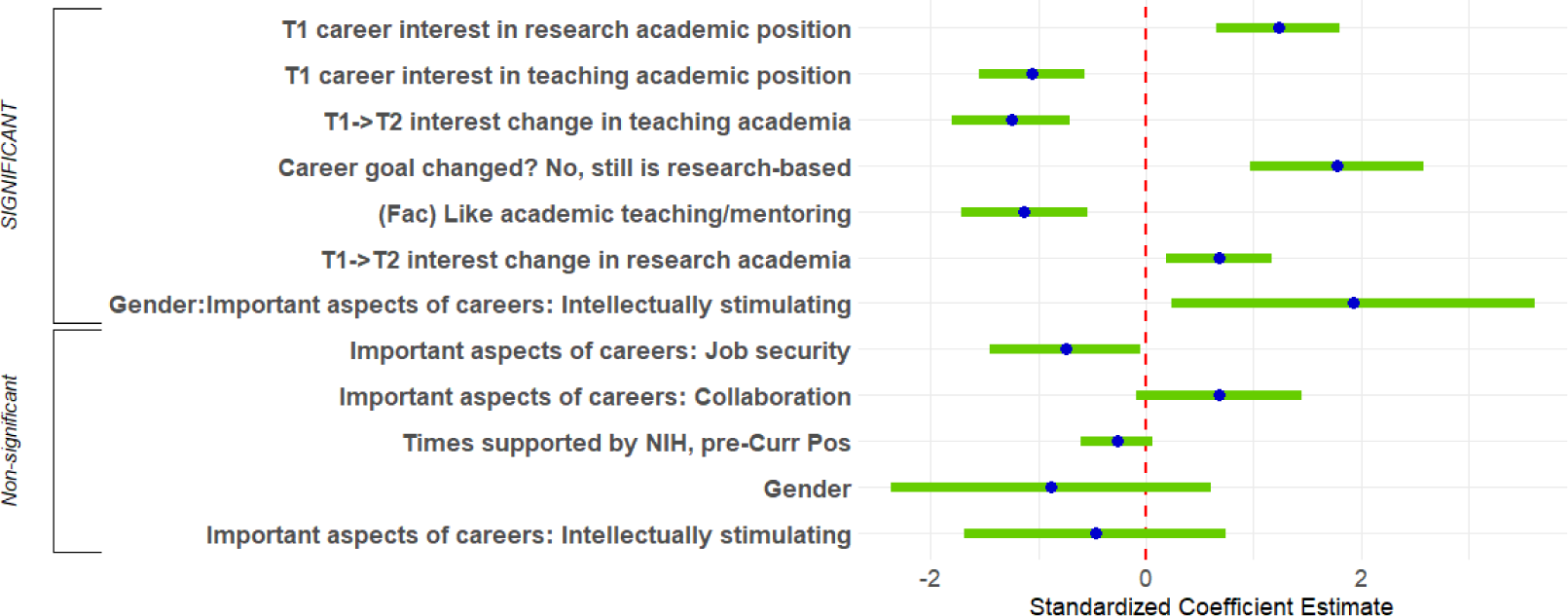
Logistic Regression Predicting Research-Focused vs. Teaching-Focused Academic Positions. Standardized regression coefficients and error bars for logistic regression predicting whether respondents were in research-focused academic positions or teaching-focused academic positions. Dependent variable was binary indicator of whether respondents’ positions were research-focused. Independent variables included career interest and change in career interest, personal characteristics, objective measures, NIH support, and interactions with gender. The entire equation was significant at p<0.001 and accurately predicted 90% of respondents in the analysis. See Extended Data Figure 5-1 for the list of explanatory/independent variables.

**Table 4.** Logistic regression predicting Research-focused vs. Teaching-focused Academia.

| Independent Variables | Unstd. Coeff. | Adj. Sig. | Odds Ratio | Effect Size | Inverse Odds Ratio |
| --- | --- | --- | --- | --- | --- |
| (Intercept) | -0.11 |  | 0.90 | - |  |
| T1 career interest in research academic | 1.23 | *** | 3.41 | (M) |  |
| T1 career interest in teaching academic | -1.06 | *** | 0.35 | (M) | 2.86 |
| T1->T2 interest change in research academia | 0.68 | * | 1.97 | (S) |  |
| T1->T2 interest change in teaching academia | -1.25 | *** | 0.29 | (M) | 3.45 |
| Important aspects of careers: Collaboration | 0.68 |  | 1.97 | (S) |  |
| Important aspects of careers: Job security | -0.75 |  | 0.47 | (S) |  |
| (Fac) Like academic teaching/mentoring | -1.13 | *** | 0.32 | (M) | 3.13 |
| Career goal changed? No, still is research-based | 1.78 | *** | 5.93 | (L) |  |
| Times supported by NIH, pre-Curr Pos | -0.27 |  | 0.76 | - |  |
| Gender | -0.88 |  | 0.41 | (S) |  |
| Aspects of careers: Intellectually stimulating | -0.47 |  | 0.62 | (S) |  |
| Gender: Aspects of careers: Intellectually stimulating | 1.93 | * | 6.90 | (L) |  |

| FULL EQUATION STATISTICS |  |
| --- | --- |
| N | 497 |
| Accuracy | 90% |
| AIC | 274.75 |
| Likelihood Ratio Test |  |
| ChiSq | 253.05 |
| p value | 0.0000 |

(Caption) Results of logistic regression predicting whether respondents were in research-focused academic positions or teaching-focused academic positions. (Fac) = variable is a result of factor analysis; Unstd. Coeff. = Unstandardized coefficient; Adj. sig. = FDR-adjusted significance. Effect sizes are labeled when they reach at least “small” size. (S) = small effect size, (M) = medium effect size, (L) = large effect size. \* $p < 0.05$ ; \*\*\* $p < 0.001$ .

Most of the significant predictors were strongly associated with being in research-focused academic positions and were either indicators of interest or personal preferences. Perhaps unsurprisingly, interest in research-focused academic positions at the start of graduate training was positively associated with being in research-focused academic positions, and interest in teaching-focused academic positions at the start of graduate training was positively associated with being in teaching-focused academic positions. In addition, increases in interest in research-focused academic positions over the course of graduate training were positively associated with being in research-focused academic positions, and increases in interest in teaching-focused academic positions over the course of graduate training were positively associated with being in teaching-focused academic positions.

In terms of personal preferences, choosing teaching/mentoring as a positive feature of academia was positively associated with being in a teaching-focused position, with a medium effect size. Similarly, indicating that one’s “career goal was research-based and had not changed” was strongly positively associated with being in a research-focused position (large effect size).

Finally, men who chose intellectually stimulating work as an important aspect of their careers were significantly more likely than women to be in research-focused positions (89%) rather than teaching-focused positions (11%) (χ^2^(4)=13.97, p<0.0001). For women, the association was not significant (an interaction).

#### Factors Associated with Being in Non-Academic Research vs. Science-Related Non-Research

Finally, we examined the associations between explanatory variables and respondents’ type of non-academic position: non-academic research or science-related non-research. The first step lasso logistic regression investigating factors associated with being in non-academic research positions vs. science-related non-research positions. Extended Data Figure 6-1 provides the list of explanatory/independent variables that were entered into the second step OLS logistic regression. The second step logistic regression equation (Table 5) was performed on data from the 284 participants who were not in academia. The equation had a significant log-likelihood test, predicting whether 84% of participants were in non-academic research positions or science-related non-research positions. Only two predictors were significant and had sufficient effect size to report. As might be expected, being in non-academic research was positively associated with both an increase in interest in non-academic research over the course of graduate training, and strongly associated with indicating that their career goal was research-based and had not changed.

**Figure 6.**
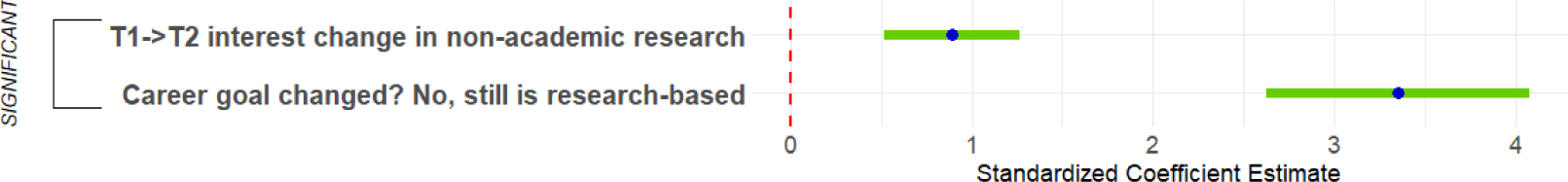
Logistic Regression Predicting Non-Academic Research vs. Science-Related Non-Research. Standardized regression coefficients and error bars for logistic regression predicting whether respondents were in non-academic research positions or scientific non-research positions. Dependent variable was binary indicator of whether respondents’ were in non-academic research positions. Independent variables included change in career interest and whether respondents’ career goals had changed. The entire equation was significant at p<0.001 and accurately predicted 84% of respondents in the analysis. See Extended Data Figure 6-1 for the list of explanatory/independent variables.

**Table 5.** Logistic regression predicting Non-academic Research vs. Science/Non-research.

| Independent Variables | Unstd. Coeff. | Adj. Sig. | Odds Ratio | Effect Size | Inverse Odds Ratio |
| --- | --- | --- | --- | --- | --- |
| (Intercept) | -1.66 | *** | 0.19 | (L) | 5.26 |
| T1->T2 interest change in non-academic research | 0.89 | *** | 2.44 | (S) |  |
| Career goal changed? No, still is research-based | 3.35 | *** | 28.37 | (L) |  |

| FULL EQUATION STATISTICS |  |
| --- | --- |
| N | 284 |
| Accuracy | 84% |
| AIC | 240.32 |
| Likelihood Ratio Test |  |
| ChiSq | 158.48 |
| p value | 0.0000 |

(Caption) Results of logistic regression predicting whether respondents were in non-academic research positions or scientific non-research positions. Unstd. Coeff. = Unstandardized coefficient; Adj. sig. = FDR-adjusted significance. Effect sizes are labeled when they reach at least “small” size. (S) = small effect size, (L) = large effect size. \*\*\* $p < 0.001$ .

### Differences in How Participants Found Their Current Positions

The final analysis investigated whether there were any differences in the primary way participants found their current positions by type of current position. An omnibus chi-squared test of independence on the contingency table was performed first and was significant (Table 6). Follow-up analyses were performed comparing types of current position for each method of finding current positions.

**Table 6.** Contingency Table Analysis of Current Position by How Participants Found Current Position.

| What was the primary way you found your current position? | Current Position | | | | | | | | TOTAL (for row) | | Sig. of Follow-up $\chi^2$ |
| --- | --- | --- | --- | --- | --- | --- | --- | --- | --- | --- | --- |
|  | Research-focused Academic |  | Teaching-focused Academic |  | Non-academic Research |  | Science Non-research |  |  |  |  |
|  | n | %Col | n | %Col | n | %Col | n | %Col | n | %Col |  |
| Contacted by employer/recruiter | 13 | 3% | 1 | 1% | 10 | 8% | 8 | 5% | 32 | 4% | n.s. |
| Former advisor/supervisor | 61<sup>1</sup> | 15% | 2<sup>2</sup> | 2% | 9<sup>2</sup> | 7% | 7<sup>2</sup> | 5% | 79 | 10% | *** |
| Job posting | 130<sup>1</sup> | 33% | 64<sup>2</sup> | 63% | 45<sup>1</sup> | 34% | 53<sup>1</sup> | 35% | 292 | 37% | *** |
| Previous position at same organization | 83<sup>1</sup> | 21% | 5<sup>2</sup> | 5% | 6<sup>2</sup> | 5% | 9<sup>2</sup> | 6% | 103 | 13% | *** |
| Professional networking (not advisor) | 76<sup>1</sup> | 19% | 28<sup>3</sup> | 28% | 56<sup>2</sup> | 42% | 57<sup>2</sup> | 38% | 217 | 28% | *** |
| Other | 33 | 8% | 1 | 1% | 8 | 6% | 16 | 11% | 58 | 7% | not performed |
| TOTAL (for column) | 396 | 100% | 101 | 100% | 134 | 100% | 150 | 100% | 781 | 100% | *** |

| FULL TABLE STATISTICS |  |
| --- | --- |
| Monte Carlo simulated chi-squared test of independence (2000 runs) | 123.01 |
| Simulated p value | 0.0005 |

(Caption) Results of contingency table analysis relating respondents' current positions and how they found their current positions. Rows show the distribution of people in current positions who found their positions through the primary method listed. The total (for row) column provides the overall distribution of ways participants found their current position. Superscripts show row groups that had similar (not-significantly different) rates. %Col = Column percentages; Sig. of Follow-up $\chi^2$ = FDR-adjusted significance of $\chi^2$ for that row. \*\*\* p < 0.001. Superscripts represent groups (by row) that are significantly different at an FDR-corrected p value of 0.05 or lower.

Participants who had found their current positions through an advisor or former advisor were more likely to be in research-focused academic positions, though it was still a relatively less common way to find an academic position. Cell-by-cell follow-up comparisons showed that the percentage of time that this occurred (15%) was significantly larger than the percentage of time for all three other types of current position, and that those three did not significantly differ from each other. A similar pattern occurred for participants who had found their current positions through a previous position at the same organization: participants in research-focused academic positions reported this avenue to their current positions at a higher rate (21%) than participants in all three other current positions, whose rates did not significantly differ from each other.

Another interesting finding concerned participants who had found their current positions through job postings. Although this was by far the most popular route to current positions (37% overall), it was significantly more prevalent for participants who were in teaching-focused academic positions (63%) than the three other types of current positions, whose rates did not significantly differ from each other.

Finally, those participants who had found their current positions through (non-advisor) professional networking were more likely to be in non-academic research positions (42%) or science-related non-research positions (38%; rates that did not significantly differ from each other) than participants in the other two current positions, whose rates did not significantly differ from each other. Research-focused academic positions were the least likely to have been found in this manner (19%, significantly lower than all other current position types).

## DISCUSSION

This work was designed to retrospectively investigate the factors that influence the career sector in which neuroscience PhDs are employed, including career interests, social identity, experiences in graduate school and postdoctoral training, and personal characteristics.

This study found that the most potent predictor of career type was an individual’s feelings about monetary compensation. This suggests that financial considerations play a crucial role in career decision-making for neuroscientists. It was one of several differentiators of academic vs. non-academic careers that were individual based: interests, values, and experiences. These factors reflect the personal motivations and aspirations of individuals and highlight the importance of aligning one’s career with their passions and values.

Our results also point to the important role that advisors and networks had on our respondents’ career development, especially during the postdoctoral phase. Advisors not only provided valuable support, but also helped their advisees secure research academic positions. Even for advisees who left academia, many of whose interests shifted gradually away from academia, advisors gave career advice that they found helpful. This underscores the significance of mentorship and guidance in shaping the career trajectories of early-career scientists. Advisors can play a vital role in helping their mentees navigate the academic landscape and make informed career choices. Although no interactions with representation status remained significant in this analysis, previous research shows that science identity and feelings of belongingness are also important for the persistence of scientists from UR groups (Fisher et al. 2019; Margherio et al. 2016; Estrada et al. 2011).

### Shifting Career Interests

We found that, in general, although most participants reported that they were interested in research-focused academic positions at the start of their PhD training, by the end of their PhD training those who would end up in different career types reported that their interests had shifted to their current career type. Only those currently in academic research positions reported sustained interest in that field. These results are consistent with other research indicating academic research is viewed as the default path, even by students at the start of their PhD (Gaughan & Robin, 2004; Helbing et al., 1998; Puljak & Sharif, 2009).

The continual shift in interest away from academia for those who ended up in non-academic positions, however, indicates that respondents felt that their increased interest in non-academic careers was not an acute event but a result of experiences, personal preferences, and mentoring up through the current time. This finding underscores the importance of mentorship and guidance for PhD students, as well as the need for exposure to a variety of career options beyond academia. It also highlights that career development is a dynamic and evolving process, and individuals may discover new interests and opportunities as they progress through their training.

### Differentiators of Current Position

Although many studies have investigated career interests among graduate students and postdoctoral fellows in STEM, few have information on the career sector in which respondents ultimately ended up working in. This study investigated which factors were associated with the type of position that respondents were in at the time of response.

Our approach was to organize our analyses as a series of binary questions that reflected the framing evident in qualitative analysis of free-text responses to this survey (Ebrahimi et al., 2022). We first considered, as many respondents may have, the broad choice of academia vs. non-academia, and asked how participants in these two sectors differed from each other. Our findings showed that participants in academic positions (both research and teaching) valued job security and autonomy and did not prioritize monetary compensation or diverse work, compared to those in non-academic positions. Those in academia were more positive about structural aspects of academia (job market, promotion, and tenure) and teaching/mentoring.

While many of these aspects seem inherent to an academic position, in fact they may not be, but more a consequence of tradition. In addition, these aspects have disparate consequences depending on an individual’s circumstance. In particular, not prioritizing monetary compensation may be a luxury that not all can afford. UR respondents selected “monetary compensation” as an important aspect of career choice more often than WR respondents and our UR respondents were twice as likely as WR respondents to have been the first person or in the first generation of their family to graduate from a 4-year college or university. Levels of student loan debt and other circumstances (such as providing financial support to extended family) vary by social identity, with UR students carrying higher levels of debt (Niu, 2016; Webber & Burns, 2022; Zeiser et al., 2013). Programs such as the NIH Loan Repayment Program attempt to retain scientists in research careers through debt reduction (National Institutes of Health, n.d.).

Valuing autonomy within academia may also have disparate impacts. As a concept, autonomy is linked to an American cultural ideal in which independent, hardworking entrepreneurship leads to wealth, status, and power. Women, lower-class men, and members of UR groups face penalties when they don’t appear to fulfill this model (Blair-Loy & Cech, 2022; Blair-Loy, 2013). If academic institutions wish to retain a more diverse faculty, they may wish to rethink incentive structures to encourage and recognize team science and collaboration (Cline et al., 2020).

An unspoken underlying issue for the next several findings is that, traditionally, funders, academic institutions, and faculty often consider student and postdoctoral “success” to equate to remaining in academia (Sauermann & Roach, 2012). While this attitude has been changing, institutions within the STEM ecosystem should continue to promote the broad array of careers for students and postdocs, and tailor training and experiences to the career path that each individual prefers (Cynthia N Fuhrmann, 2016; Lenzi et al., 2020). Our findings suggest that personal history, interests, and values are all factors that contribute to an individual’s career interests and outcome.

Faculty involvement in our academic respondents’ histories was a complicated picture. Faculty support during their postdoctoral training was strongly associated with remaining in academia. Respondents’ ratings of helpfulness of postdoctoral advisor career advice, however, were negatively associated with remaining in academia. It is possible that career advice as a postdoctoral fellow was more important for those who were thinking about leaving academia than those who wanted to pursue academic careers. Since postdocs as a group will generally contain both those who wish to remain in academia and those who wish to pursue other careers, academic institutions should consider fostering both faculty support of postdocs and career advising to address both groups.

Social identity was also associated with differences in respondents’ histories. For women, but not men, feelings of departmental belongingness during postdoctoral training were significantly positively associated with remaining in academia. Sense of belonging has been shown to be related to persistence in science, especially for underrepresented groups (Good et al., 2012; Luft et al., 2004; Yen et al., 2017). The postdoctoral experience can be especially isolating, but institutions can combat this by establishing or supporting Offices of Postdoctoral Affairs and postdoctoral associations (Åkerlind, 2005; Fork et al., 2020; Graham et al., 2018; Nowell et al., 2018).

Our second question was, if respondents remained in academia, what factors differentiated those who were in primarily research-based positions from those who were in primarily teaching-based positions? The answers were clear and perhaps unsurprising: the strongest differentiating factors were early interest (at the start of graduate school) in each type of position, becoming more interested in each type of position over the course of graduate school, and initial and sustained interest in research, which was associated with being in a research-focused position.

A more surprising finding was that choosing teaching/mentoring as a positive feature of academia was negatively associated with being in a research-focused academic position. This might not have been expected to differentiate as strongly, considering that teaching and mentorship are often considered to be a key aspect of all academic faculty positions. This aspect of the job, however, may not be as important to those who are drawn to research-focused positions. Given the interdependency between quality training and research accomplishment, these data may highlight the critical need for focused and specific mentor training built into all environments to ensure teaching and mentorship remains valued.

In addition, men, but not women, who chose “intellectually stimulating work” as an important aspect of their careers were significantly more likely to be in research-focused positions. This may indicate that women have a broader definition of what they consider “intellectually stimulating” work than men.

Our final question was, if respondents were not in academic faculty positions, what factors differentiated those who were in non-academic research positions from those who were in non-research scientific positions? The answers to this question were straightforward—a strong and/or enduring interest in research. Being in non-academic research was positively associated with both an increase in interest in non-academic research over the course of graduate training and strongly associated with indicating that their career goal was research-based and had not changed. No other factors differentiated these two professions; perhaps because the comparison between them was not reflective of actual choices that people might have made. Although the decision to stay in academia may be an overarching question for students and postdoctoral fellows, what to do if one leaves academic may not be as clear cut a choice. Choosing a non-academic career may depend on different variables than those that we measured, and it may depend on many variables, not just a few.

Interestingly, however, the analyses discussed above included only a few interactions with gender, and no interactions with representation status. This may indicate that although there are gender and UR differences in the proportion of respondents in the different career types, the factors that led to these differences were less about how the same factors affected social identity groups differently (interactions in prediction), but that different social identity groups had different interests, values, and experiences that led to different careers.

### How Participants Found Their Current Positions

There were clear differences in how respondents in different types of positions found their jobs. On the one hand, those in research-focused academic positions were the most likely to have found their current positions through an advisor or former advisor or found their current positions through a previous position at the same organization. Indirect (advisors, networking) or direct (previous positions) familiarity seems to have been an important factor in hiring decisions for these positions (see similar findings among NINDS Early Stage Investigator R01 applicants in Hsu et al. (2021)). Similarly, professional networking was the predominant method for those in non-academic research positions or science/non-research positions. On the other hand, job postings, perhaps considered the default way to find a job, was the much more prevalent method for participants who were in teaching-focused academic positions—almost double the rates of the other three types of careers.

These findings underscore the role of networking and personal relationships in securing employment in STEM. Reliance on these informal and formal relationships are likely to disadvantage women and underrepresented scientists. Research has shown that women have less access to sponsorship, support, and mentoring opportunities (Moss-Racusin et al., 2012; Nolan et al., 2008; Paksi & Tardos, 2021; Patton et al., 2017) and are less likely to benefit from networking for career-enhancement (Forret & Dougherty, 2004; Gersick et al., 2000). In fact, we found that women were approached by employers or recruiters for their current positions less often than men were.

Members of underrepresented groups also have less access to social capital through differences in size and composition of networks and strength of ties within their network (Forret, 2006). For example, Pinheiro & Melkers (2011) found that UR scientists have a greater proportion of collaborative ties outside their home institutions, but this was related to a lack of support at the home institution and chilly work climate. In Ullrich et al. (2021), we also found that UR respondents reported more beneficial relationships with faculty outside their PhD institutions than WR respondents. Projects within the NIH Neuroscience Development for Advancing the Careers of a Diverse Research Workforce R25 program support mentoring and scientific networks at a national level to address some of these factors that can isolate UR scientists and hinder their career progress (National Institute of Neurological Disorders and Stroke, 2023).

### Limitations

This study has the same limitations as discussed in Ullrich et al. (2021), which reported on respondents to the same survey: it is limited to citizens and permanent residents who have applied or been appointed to NINDS grants; it is not a random sample of respondents; it has a large number of explanatory variables; it does not reflect the effects of the COVID-19 pandemic; and it relies on retrospective reports. Due to the nature of the regression methods used, it also collapses rich, multifaceted social identities into large enough sample sizes to analyze, namely binary gender and binary groups of race/ethnicity (WR and UR respondents). In addition, despite collapsing respondents into a single UR group, it is possible our sample size was still too small to capture any but the largest differences by representation status.

Although our sample limitations restrain the breadth of the conclusions that we can draw, they are not outliers in the literature, since we have a similar sample demographic as other studies (Gibbs et al., 2014). As noted in the methods section, we have tried to reduce the impact of the large set of explanatory variables statistically by utilizing penalized regression techniques.

Relatedly, these findings are retrospective, and do not constitute independent data points. Unlike a prospective study, which would allow us to understand participants views at the time, retrospective ratings reflect participants’ current understanding of their interest in the past and better represent their personal narrative of their career trajectories.

A different type of limitation involves broader factors that are hard to quantify and account for. It is important to remember that although this survey is framed in terms of individual preferences and experiences, the discourse of “personal choice” can obscure the systemic pressures that different groups face (Beddoes & Pawley, 2014). For example, for women doctorates in STEM, having children and being in a dual-earner marriage are strongly associated with lower rates of labor force participation and higher rates of part-time work, likely due to women taking on the burden of family and household responsibilities (Dever et al. 2008; Waaijer et al. 2016; Bloch et al. 2015; Frank 2019; Shauman 2017; Schiebinger and Gilmartin 2010).

Other factors, such as biases in hiring decisions, may also affect employment. Biases against high-performing women or assumptions around relationship or (future) parental status can introduce gender bias during hiring that may influence patterns of workforce distribution (Moss-Racusin et al., 2012; Quadlin, 2018; Rivera, 2017; Sheltzer & Smith, 2014; Way et al., 2016). Similarly, cognitive biases against UR groups have resulted in lower call-back rates during hiring (Bertrand & Mullainathan, 2003; Eaton et al., 2019; but see Lee & Savoy, 2015). These factors, although potentially strong influences on employment outcomes, are impossible to capture in studies of employees (rather than studies of employers) due to survivorship bias.

## Conclusion

To ensure that all neuroscientists, and especially underrepresented or women neuroscientists, are not driven away from academic research, but instead feel free to follow their passions (whether in academia or not), institutions and advisors will need to play a key role in reframing academic culture to value teaching, mentoring, and inclusive environments and better preparing graduate students and postdoctoral fellows for the career outcomes that match their preferences (see Ullrich et al., 2021 for discussion of potential interventions in this space). To create a more inclusive and diverse academic environment, it’s essential to address financial disparities, provide tailored support, foster mentorship relationships, and actively work to create inclusive academic cultures that embrace a variety of career paths for neuroscientists.

## ACKNOWLEDGMENTS

We thank the Diversity Working Group at NINDS for feedback and input (especially Edgardo Falcon-Morales, Katie Pahigiannis, Ashlee Van’t Veer, and Letitia Weigand for initial survey design). We also thank Thomas Cheever, Devon Crawford, Anahid Ebrahimi, Jordan Gladman, Mariah Hoye, Jenny Kim, Marguerite Matthews, Marilyn Moore-Hoon, and Ling Wong for feedback on this manuscript draft. In addition, we thank Walter Koroshetz and Janine Clayton and the NIH Office of Research on Women’s Health for their leadership and financial support.

